# Alzheimer’s disease-associated P460L mutation in ephrin receptor type A1 (EphA1) leads to dysregulated Rho-GTPase signaling

**DOI:** 10.1101/2021.06.17.448790

**Authors:** Y. Kim, G. Lasso, H. Patel, B. Vardarajan, I Santa-Maria, R. Lefort

## Abstract

Recently, late onset AD (LOAD) genome-wide association studies identified EphA1, a member of receptor tyrosine kinase family (RTK) as a disease associated loci. In the follow-up study where 3 independent LOAD cohorts were performed, a P460L coding mutation in EphA1 loci showed a significant association with LOAD. However, the role of EphA1 and P460L mutant EphA1 in AD is not fully understood. We have characterized this mutation biophysically and biochemically. Our structural in silico model and in vitro biochemical analysis demonstrate that EphA1-P460L mutation makes the receptor constitutively active suggesting a gain-of-toxic function leading to chronic EphA1 signaling in the brain. Moreover, we report that the EphA1 P460L variant triggers Rho-GTPase signaling dysregulation that could potentially contribute to spine morphology abnormalities and synaptic dysfunction observed in AD pathology.

## INTRODUCTION

Large-scale genome-wide association studies (GWAS) and whole genome and exome sequencing have identified several variations in the human genome that significantly affect susceptibility to late-onset Alzheimer’s disease (LOAD). While these genes collectively implicate several pathways (e.g. synaptic function, inflammation, lipid metabolism, endocytosis/intracellular trafficking, and axonal transport) in LOAD pathogenesis, evidence supporting altered function in specific genes and more particularly variant-specific expression and effect on LOAD etiology for most of these genes remains incomplete. We recently published the results of targeted sequencing of *ABCA7, BIN1, CD2AP, CLU, CR1, EPHA1, MS4A4A/MS4A6A, and PICALM* in three independent LOAD cohorts (Caribbean Hispanic families, NIA LOAD family studies and unrelated Canadian individuals of European ancestry), where we found several non-synonymous mutations in all three cohorts, including in EPHA1 (P460L), which segregated completely in an extended Caribbean Hispanic family [1].

EPHA1 encodes for the Eph receptor A1, a member of the largest family of receptor tyrosine kinases (RTK), the erythropoietin-producing hepatocellular (Eph) family [2, 3]. Eph receptors (EphRs) are divided into 2 subclasses (EphA and EphB), comprised of several members (EphA1-A10 and EphB1-B6), which interact with various ephrin ligands (ephrin-A1-A5 and ephrin-B1-B3) [4]. EphRs and ephrin ligands can signal independently of each other, and/or in concert with other cell-surface receptors. For example, members of the epidermal growth factor (EGF) receptor family have been shown to bind EphA2 to promote cell migration and proliferation [5], while members of the fibroblast growth factor (FGF) receptors have been demonstrated to form signaling clusters with EphA4 to regulate cell proliferation [6-8]. The vast number of interacting partners and signaling methods allow EphRs and ephrin ligands to control numerous developmental processes and to regulate the normal physiology and homeostasis of many organs, including the central nervous system (CNS) [9]. Conversely, aberrant Eph/ephrin signaling has been implicated in a variety of diseases, from cancers [10, 11] to neurological diseases, such as Amyotrophic Lateral Sclerosis (ALS) [12]. To date, in addition to EphA1, two other EphRs have been implicated in LOAD, EphA4 and EphB2 [1, 13-19], which means that of the sixteen known EphRs, three are implicated in LOAD. This is not entirely surprising given the number of complex signaling networks in which EphRs and their ligands participate. One intriguing possibility is that disruption at a single node of this network, due to a function-altering mutation for example, is sufficient to completely disturb cellular or circuit homeostasis in the brain. We set out to determine which pathways might be disrupted by the AD-associated P460L variant of EphA1.

We report here that the EphA1-P460L variant results in the constitutive activation of the receptor and of its kinase activity. Computational modeling and experimental analysis reveal that the substitution alters the receptor’s interaction with the lipid bilayer, possibly restricting its lateral movement and favoring homo-interactions, receptor clustering and auto-activation. Constitutive activation of the EphA1 leads to aberrant downstream signaling of the small RhoGTPases, RhoA and Rac1, as previously demonstrated in several cellular and animal AD models [20-30].

## RESULTS

### EphA1 P460L amino acid substitution promotes the embedding of its ectodomain onto the membrane

Nonsynonymous single nucleotide polymorphisms (nsSNPs) introduce amino acid changes in their corresponding proteins potentially altering folding or complex assembly, resulting in profound effects on the enzyme activity of the protein. The magnitude of the substitution, however, is often difficult to predict. EphA1 protein structure is not yet fully known. Previous computational and experimental analyses show that the EphA2 fibronectin type III domain repeat 2 (FN2), proximal to the membrane, interacts with lipids through electrostatic interactions between an electropositive patch of FN2 and the polar head groups of anionic lipids [31]. Human EphA2 and EphA1 receptors share a sequence identity of 52% and have same domain architecture (Figure S1A). Our predicted structural model and electrostatic calculations show that EphA1 FN2 domain has an electropositive patch that structurally aligns with the lipid binding electropositive patch of EphA2-FN2 (Figure S1B), suggesting a similar lipid binding role of the EphA1 FN2 domain. P460 lies on the same face of the domain as the electropositive patch and structural superposition between the modelled EphA2 FN2-lipid complex [31] and the EphA1 FN2 atomic model suggests that P460 is partially embedded into the lipid bilayer (Figure 1B, C). Modeling and structure analysis suggest a potential role of P460L mutation in stabilizing the EphA1 ectodomain onto the lipid bilayer through enhanced hydrophobic contacts with the lipids and promoting clustering and activation of EphA1 receptors independently of ligand binding.

**Figure 1.**
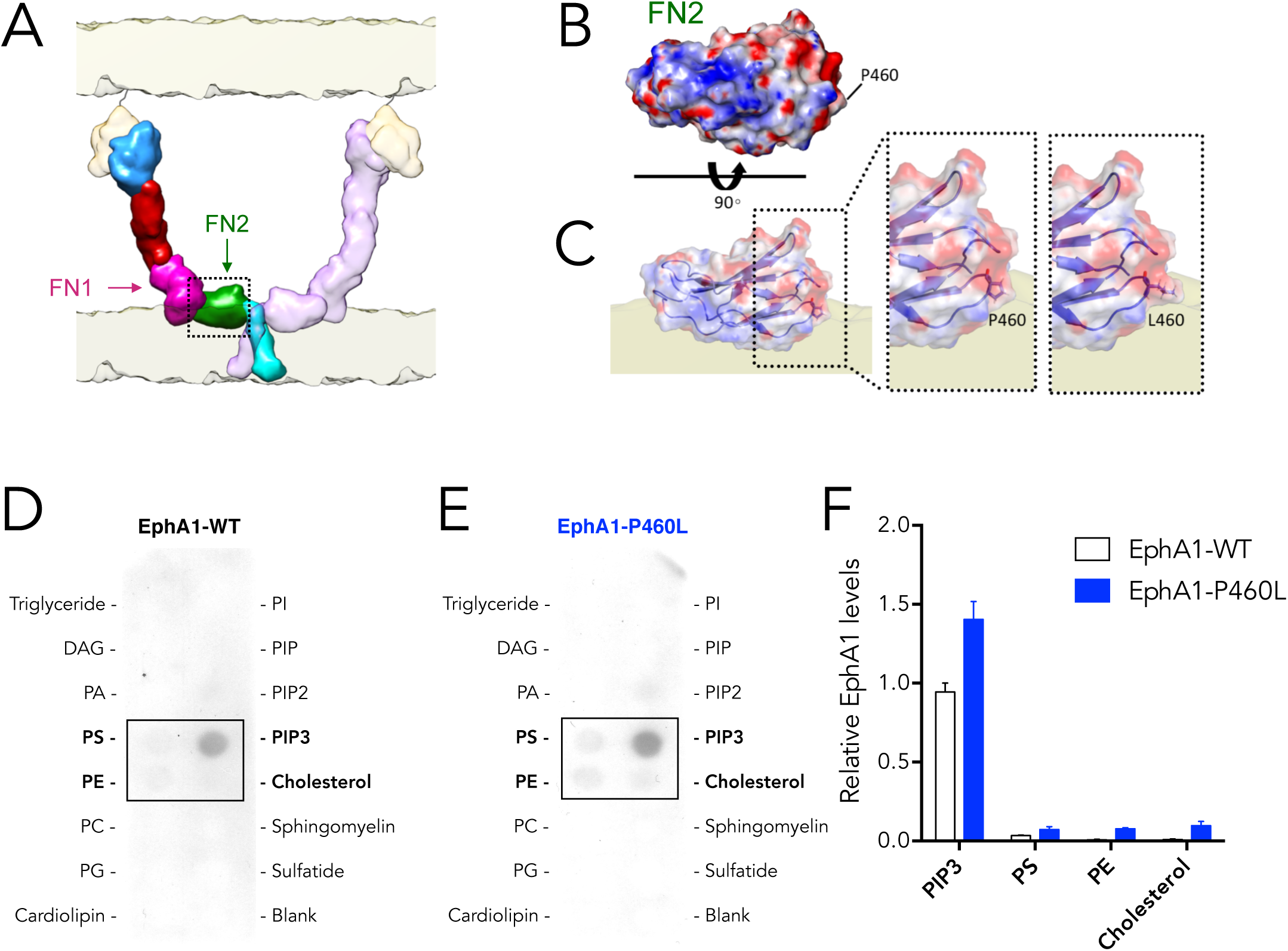
Modeling of the EphA1 FN2 domain and structural comparison with EphA2 FN2 domain. (A) Structural superposition of the EphA1 FN2 atomic model (blue) and EphA2 FN2 (red). (B) Electrostatic surface of the EphA1 FN2 atomic model. (C) Electrostatic surface of the EphA2 FN2 crystal structure. (Scale bar: red, -5kT; blue, +5kT). (D-F) Lipid binding assay showing enhanced interactions of EphA1-P460L variant with Phosphatidylinositol (3,4,5)-trisphosphate (PIP3), cholesterol and Phosphatidylethanolamine (PE). (D, E) Representative Western blot of PIP strips membranes of EphA1 interactions with several components of the lipid bilayer. (F) Quantification of Western blot data. Protein quantifications were normalized to total lysate protein concentration. Data presented as mean ± s.d. of N=3 independent experiments.

To test this hypothesis, we performed a cell-free lipid binding assay to determine the interaction of EphA1-P460L with membrane lipids. We generated stable HEK293 (293) cell lines overexpressing either wild-type EphA1 (EphA1-WT) or EphA1-P460L (Figure S2). Whole proteins were isolated from these cells under native conditions and incubated with membranes on which various components of the lipid bilayer were adsorbed. As predicted from our computational analysis, we found that EphA1-P460L interacts with a higher affinity with some lipids, specifically with phosphatidylinositol (3,4,5)-trisphosphate (PIP3), phosphatidylserine (PS), phosphatidylinositol (PE) and cholesterol (Figure 1D-F). These results are in agreement with our predicted model, suggesting that EphA1 P460L substitution may promote embedding of the EphA1 receptor into the lipid bilayer.

### EphA1 P460L substitution enhances receptor auto-activation independently of ephrin ligand stimulation

Forward signaling of Eph receptors is similar to the prototypical RTK mode of signaling, which is triggered upon ligand binding resulting in auto-activation phosphorylation and activation of the kinase domain [32]. What sets these receptors apart, however, is that the interaction of the ephrin ligands on juxtaposed cell surfaces with Eph receptors leads to rapid oligomerization through not only Eph receptor–ephrin interfaces but also receptor–receptor cis interfaces located in multiple domains resulting in large signaling clusters [33, 34], which can enlarge to incorporate Eph receptors that are not bound to ephrins [35].

Our computational and experimental analysis suggest that the EphA1 P460L substitution may hinder lateral diffusion of the receptor. If correct, we would predict that EphA1-P460L receptors might be more likely to exist in large stable clusters, independently of ephrin ligand-binding thereby reducing the pool of EphA1-P460L monomers at the cell surface, relative to EphA1-WT. To confirm this prediction, the HEK293 cells were transiently transfected with either EphA1-WT or EphA1-P460L, both tagged with Myc and Flag sequences (Figure 2A). After 48 hours the cells were incubated with the membrane-impermeable, cleavable cross-linker DTTSP (3,3’-dithiobis(sulfosuccinimidyl propionate)) to cross-link cell surface proteins (Figure 2B). Cross-linked EphA1 clusters were subsequently isolated from lysates by immunoprecipitation with a Myc antibody and resolved under native conditions and analyzed by Western bot using a Flag antibody. As predicted, we found a larger of pool high molecular weight clusters of EphA1-P460L relative to EphA1-WT (Figure 1C lanes 1 vs 2, E). Conversely, we found a lower pool of EphA1-P460L monomers compared to EphA1-WT (Figure 1D lanes 1 vs 2, F), consistent with the fact that most EphA1-P460L receptors reside in large clusters at the cell surface. Cleavage of the cross-linker by reducing the thiol bond resulted in the breaking-up of these clusters, confirming equal pool of available EphA1 receptors for both WT and P460L in the cells (Figure 1C, lanes 3 vs 4).

**Figure 2.**
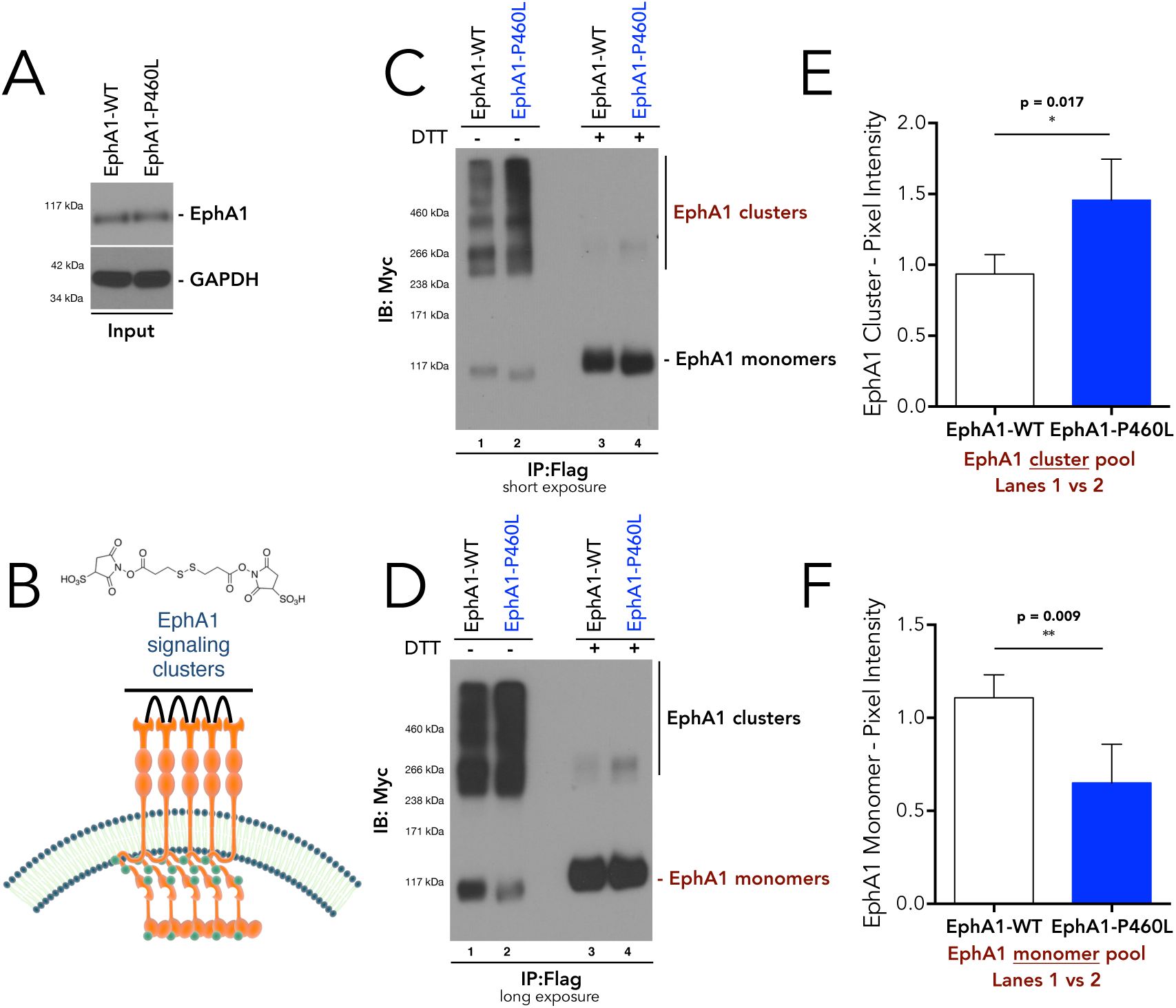
EphA1 P460L amino acid substitution promotes receptor clustering. (A) Representative Western blot showing expression of EphA1-WT and EphA1-P460L in transfected HEK293 cells. (C-D) Amino acid substitution P460L in EphA1 results in enhanced receptor clustering: Myc-Flag-tagged EphA1 (WT and P460L) was overexpressed in HEK293 cells, immunoprecipitated (Myc) and separated on a native gel to resolve all forms of EphA1; DTT was used to break up clusters. (C) Representative immunoblot (Flag) showing higher levels of high-molecular weight clusters of EphA1-P460L relative to EphA1-WT. (D) Longer exposure of immunoblot shows reduced levels of the monomeric pool of EphA1-P460L compared to EphA1-WT, consistent with higher amounts of clustered EphA1-P460L. (E-F) Densitometric quantification of (C) EphA1 clusters showing higher levels of EphA1-P460L, and (D) EphA1 monomers showing lower levels of EphA1-P460L; data presented as the mean ± s.d. (N=3 independent experiments; two-tailed unpaired t-test).

As mentioned above, assembly of Eph receptors into large clusters results in auto-phosphorylation and activation of the kinase activity of the receptor. To confirm autophosphorylation of the receptor in the absence of a ligand, transiently transfected 293 cells (EphA1 WT and P460L, Figure 3A) were collected and the immunoprecipitated proteins were immunoblotted with a phospho-tyrosine antibody (pTyr). Tyrosine-phosphorylation of EphA1 was significantly higher for P460L variant relative to WT, which showed only basal levels of tyrosine phosphorylation (Figure 3B lanes 3 vs 5, C). To confirm that both WT and P460L were still responsive the ligand binding, we incubated the transfected cells with clustered ephrin-A1 ligands for 15 minutes prior to lysing the cells. Immunoblot analysis showed significantly higher levels of phospho-tyrosinated EphA1-WT after ligand activation compared to basal conditions (Figure 3B lanes 3 vs 4, C). Interestingly, activation of EphA1-WT after ligand stimulation was similar to unstimulated EphA1-P460L (Figure 3B lanes 5 vs 6, C). In fact, ligand stimulation of EphA1-P460L only resulted in ∼60% increase in phosphorylation, suggesting that the majority of EphA1-P460L may already be phosphorylated.

**Figure 3.**
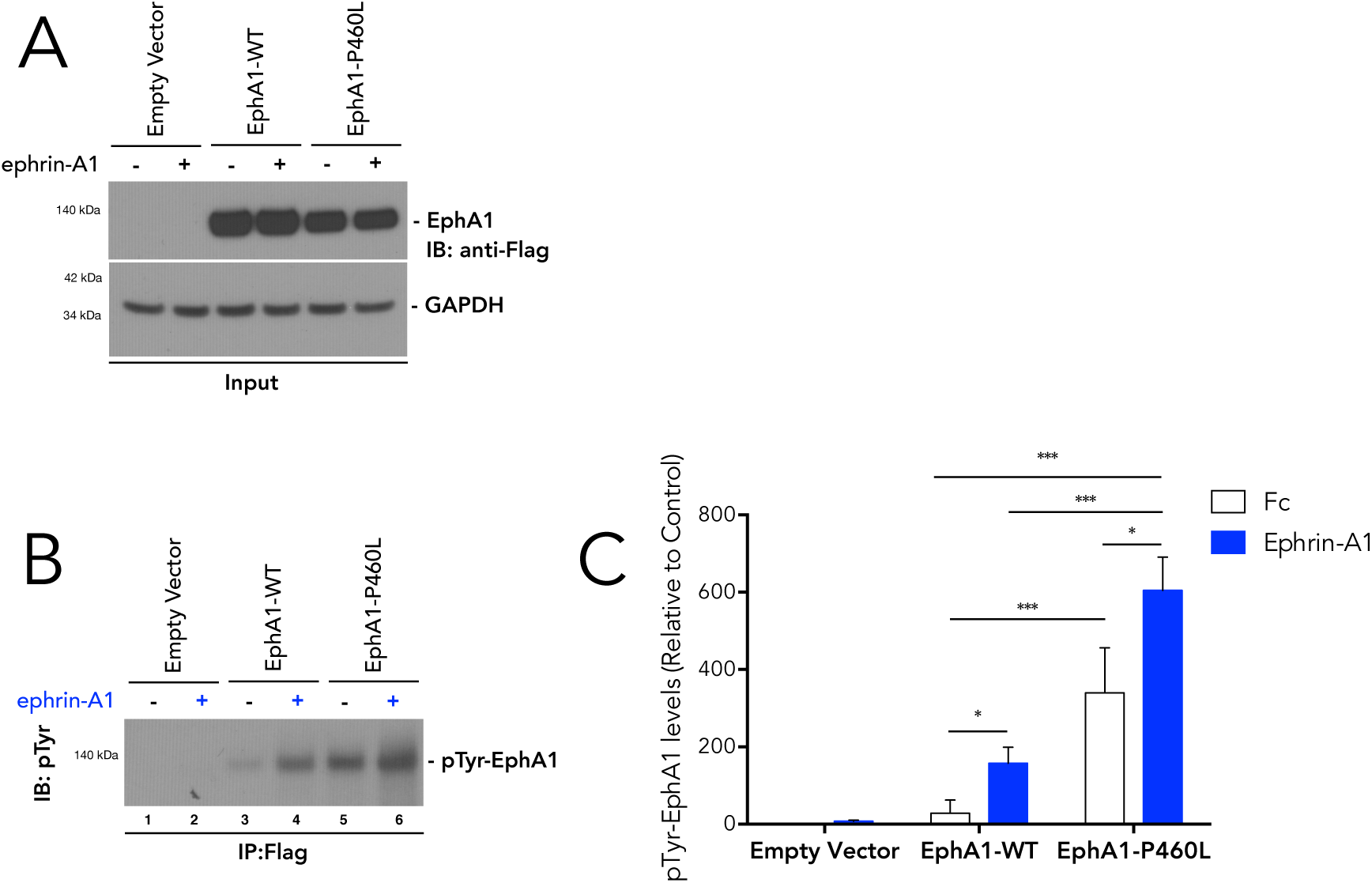
EphA1 P460L amino acid substitution results in receptor activation independently of ligand binding. Myc-Flag-tagged EphA1 (WT and P460L) was overexpressed in 293 cells, which were treated with the EphA1 ligand ephrin-A1 (10 minutes) to activate the receptor. EphA1 was immunoprecipitated (Myc) and immunoblotted with a pan phospho-tyrosine (pTyr) antibody to detect active tyrosine-phosphorylated EphA1. (A) Representative immunoblot (Flag) of input. (B) Representative immunoblot (pTyr) showing higher levels of tyrosine-phosphorylated EphA1 (pTyr-EphA1) in EphA1-P460L compared to EphA1-WT, independently ephrin-A1 stimulation. (C) Densitometric quantification of (f) pTyr-EphA1 showing higher levels of pTyr-EphA1-P460L compared to pTyr-EphA1-WT; data presented as the mean ± s.d. (N=4 independent experiments; two-way ANOVA and Bonferroni post-hoc analysis, *p<0.05, **p<0.01, ***p<0.001).

To further confirm that these clusters where active, we looked at a well-known downstream target of EphA1 signaling, namely ERK1/2. We found the levels of phosphorylated ERK1/2 to be significantly higher in EphA1-P460L-expressing cells, in a gene dose-response manner, compared to EphA1-WT cells (Figure S3), consistent with increased kinase activity. Together, these results are consistent with the idea that the P460L substitution on EphA1 results in the constitutive activation of the receptor, resulting from enhanced clustering independently of ephrin ligand binding.

### P460L substitution results in cell spreading independently of ligand stimulation

While EphA1 is the founding member of the Eph receptor family, little is known about its physiological role. It has been reported that EphA1 plays a role in cell spreading and motility [36]. Therefore, to gain insight into the functional consequences of the P460L substitution in EphA1, we decided to measure cell spreading as read-out of altered EphA1 functions. For these assays, we used the previously generated stably expressing EphA1-WT and P460L 293 cell lines (Figure S2). Cells were detached from the culture plates and seeded at equal density either on fibronectin (FN)-coated or non-coated (plastic) plates and allowed to attach and spread for 30 minutes. The cells were then stained with fluorescently-labeled phalloidin and analyzed. We found that most (∼70%) EphA1-WT expressing 293 cells plated on FN-coated plates were able to successfully attach and spread, similar to control 293 cells (Figure 4B, D, H). In contrast, significantly fewer (∼30%) of the EphA1-P460L expressing 293 cells were able to spread, retaining their circular shape (Figure 4F, H). In addition, average cell size was significantly smaller in cells expressing EphA1-P460L compared to EphA1-WT (Figure 4G, I). It should be noted that the defect appears to be a delay rather than an inability to attach and spread as most EphA1-P460L cells successfully spread after 2 hours (Figure S4).

**Figure 4.**
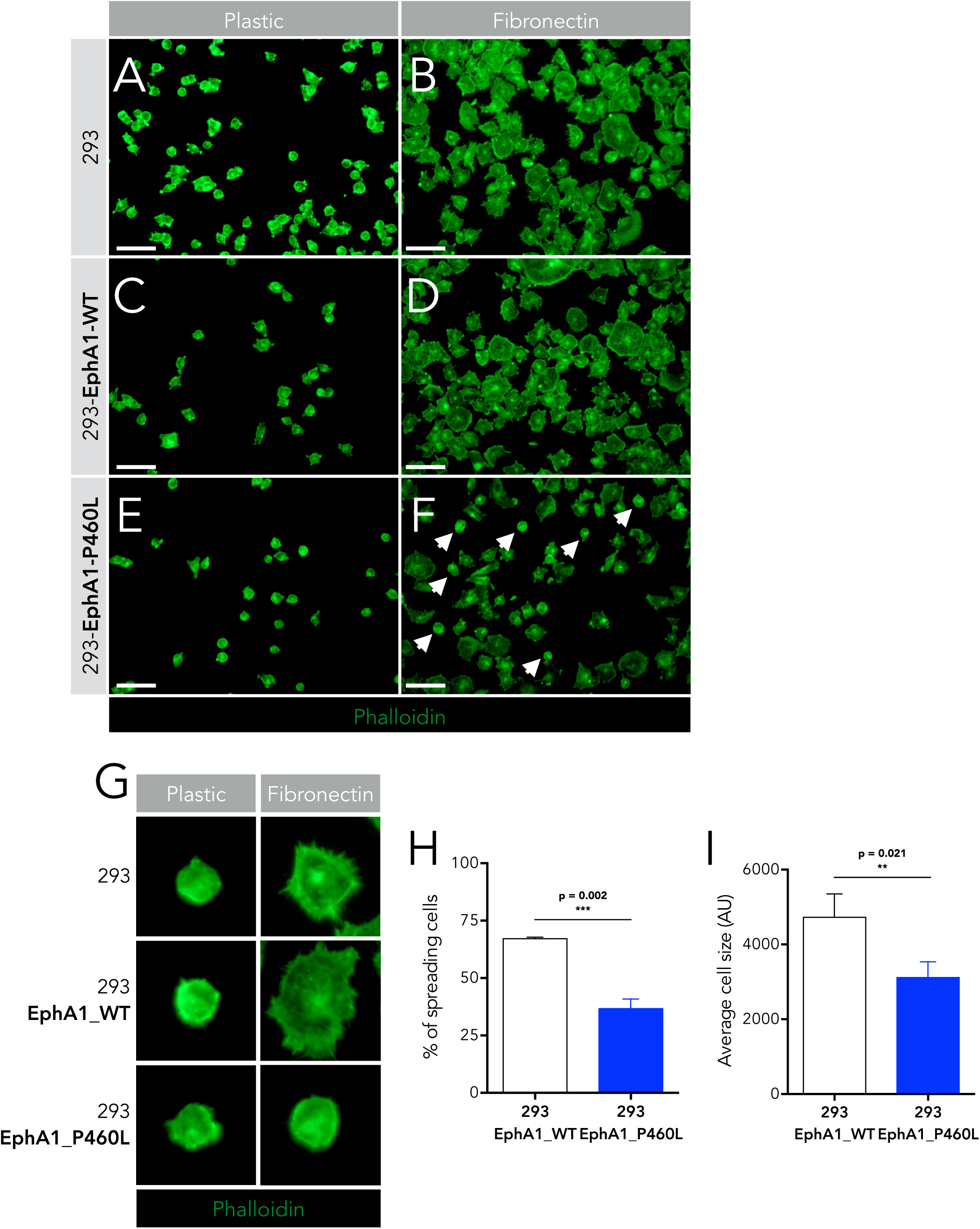
Cells expressing EphA1 P460L exhibit delayed cell spreading. (A-F) Representative images of wild-type 293 (A,B) and stably expressing EphA1-WT (C,D) or EphA1-P460L (E,F) plated on uncoated (A,C,E) or fibronectin (FN)-coated (B,D,F; 1 μg/ml) coverslips for 30 minutes and stained with Alexa Fluor-488-conjugated Phalloidin (white arrows indicate round non-spreading cells). (G) Representative images of spreading cells on FN-coated coverslips and non-spreading cells on plastic. (H) Quantification of spreading cells expressed as the percentage of the total number of cells showing a lower percentage in EphA1-P460L expressing cells compared to EphA1-WT. (I) Quantification of the average cell size showing that EphA1-P460L expressing cells are on average smaller than EphA1-WT. Data presented as the mean ± s.d. (10 randomly selected high-power fields from N=3 independent experiments; two-tailed unpaired t-test).

To confirm that the spreading defect is dependent on the kinase activity of EphA1, cells were stimulated with pre-clustered recombinant ephrin-A1 ligand (ephrin-A1–Fc; 1 μg/ml) for 15 minutes prior to detaching and replating on fibronectin-coated plates. After being allowed to spread for 30 minutes cell morphology was analyzed as above. Unlike non-stimulated cells, ligand-stimulated EphA1-WT-expressing 293 cells exhibited a significant defect in spreading (Figure 5A, B), similar to EphA1-P460L-expressing cells. Interestingly, the spreading defect phenotype was not exacerbated in EphA1-P460L-expressing after ligand stimulation, again suggesting that EphA1-P460L signaling has peaked. As an alternative method to confirm the involvement of the kinase activity of EphA1 in the spreading defect, we mutated the kinase domain of EphA1-P460L and generated stable 293 cells (EphA1^KD^-P460L. Once again, cells were detached, replated, and allowed to spread for 30 minutes before being analyzed. We found that cells expressing EphA1^KD^-P460L were able to spread normally compared to EphA1-P460L cells (Figure 5C-F). Taken together, our data shows that the P460L substitution on EphA1 alters receptor function through the constitutive activation of its kinase activity.

**Figure 5.**
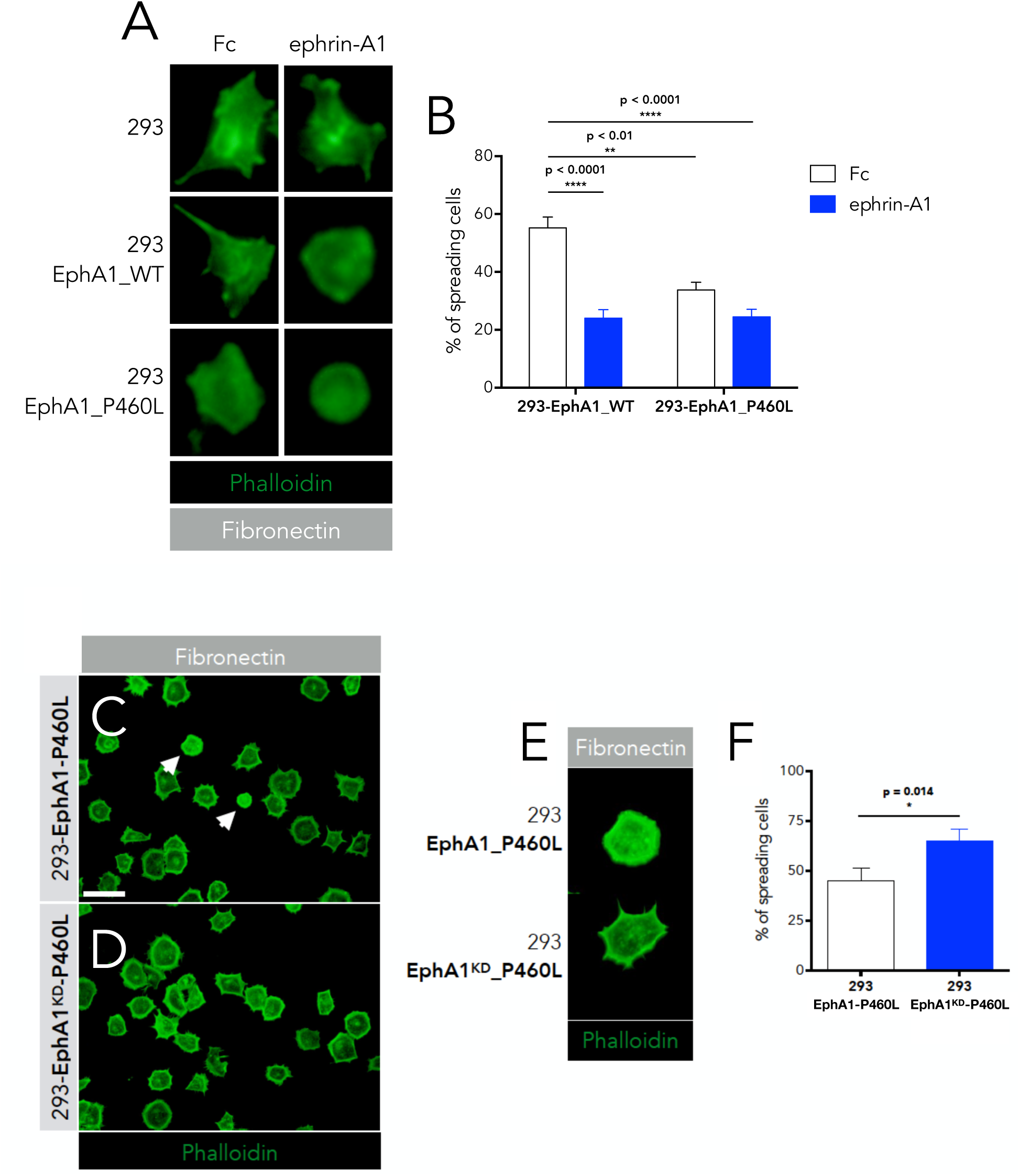
Cells expressing EphA1 P460L exhibit delayed cell spreading in a kinase-dependent manner but independently of ligand-binding. (A) Representative images of spreading cells on FN-coated coverslips and non-spreading cells on plastic. Cells were pre-treated with either pre-clustered ephrin-A1–Fc (1 μg/ml) or Fc (Control) for 15 minutes prior to plating. (B) Quantification of spreading cells expressed as the percentage of the total number of cells showing a lower percentage in EphA1-WT expressing-cells stimulated with ephrin-A1 and in EphA1-P460L expressing cells compared to control (Fc-treated) EphA1-WT expressing cells. Data presented as the mean ± s.d. (10 randomly selected high-power fields from N=3 independent experiments; two-way ANOVA followed by Bonferroni Post Hoc multiple comparisons). (C-E) Representative images of 293 cells stably expressing EphA1-P460L (C) or kinase-dead EphA1^KD^-P460L (D) plated on FN-coated (1 μg/ml) coverslips for 30 minutes and stained with Alexa Fluor-488-conjugated Phalloidin (white arrows indicate round non-spreading cells). (E) Quantification of spreading cells expressed as the percentage of the total number of cells showing a higher percentage of spreading in cell expressing kinase-dead EphA1-P460L. Data presented as the mean ± s.d. (10 randomly selected high-power fields from N=3 independent experiments; two-tailed unpaired t-test).

### Delay in cell spreading by EphA1 P460L is rescued by aberrant RhoA and Rac1 signaling

Eph receptors can regulate cell morphology through tight control of the actin cytoskeleton. A key component of this regulatory mechanism is the Rho family of small GTPases, including RhoA, Cdc42 and Rac1 [37]. These act as binary molecular switches that shuttle between an inactive (GDP-bound) and an active (GTP-bound) form in which they bind to and activate their specific downstream effectors, such as ROCK and PAK [38]. The activation of Rho GTPases is regulated by guanine nucleotide exchange factors (GEFs), which enhance the exchange of bound GDP for GTP and, thus, activate the Rho GTPases, and GTPase-activating proteins (GAPs), which inhibit Rho family members by potentiating their intrinsic GTPase activity. Direct recruitment of GEFs is one of the mechanisms by which Eph receptors mediate their functions in neurons.

Indeed, Yamazaki demonstrated that EphA1 activation affects cell spreading by altering the signaling equilibrium between the small Rho-GTPases, RhoA and Rac1 [36]. To determine whether the P460L substitution alters RhoA and Rac1 activity, we compared the levels of active RhoA and Rac1 in non-stimulated and ligand-stimulated stably expressing either EphA1-WT or EphA1-P460L. We observed a significant decrease in active Rac1-GTP levels in EphA1-P460L-expressing cells under basal (non-stimulated) conditions relative to EphA1-WT-expressing cells (Figure 6A, B). Ligand stimulation of the cells with clustered ephrin-A1 resulted in a significant attenuation Rac1 signaling in EphA1-WT-expressing cells, consistent with the requirement of kinase activation of the receptor. However, we did not observe any further dampening of the Rac1 activity in EphA1-P460L-expressing cells after ligand stimulation.

**Figure 6.**
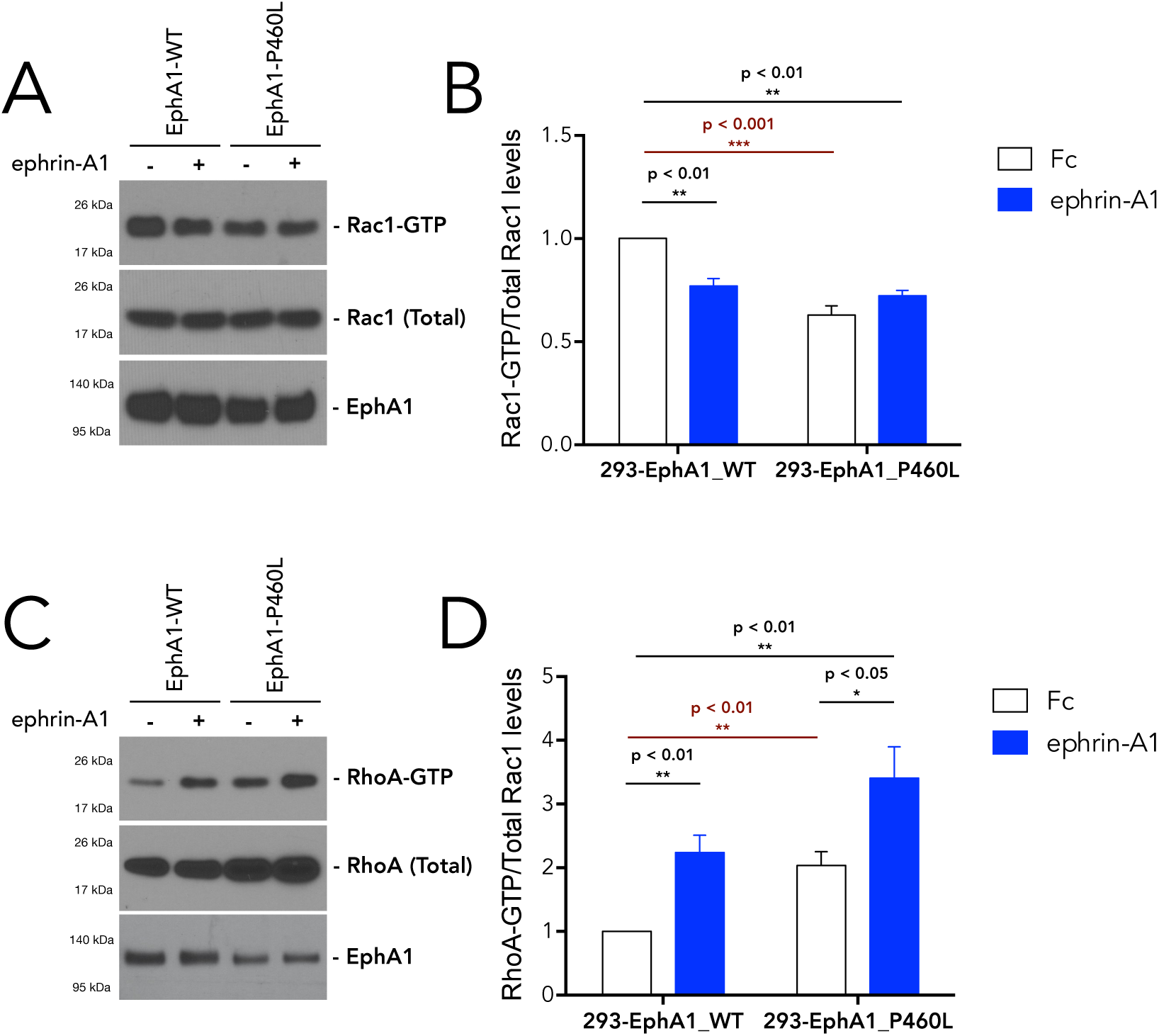
Cells expressing EphA1 P460L exhibit dysregulated RhoA and Rac1 signaling in a ligand independent manner. (A,C) Representative Western blots showing levels of active RhoA-GTP and Rac1-GTP and total RhoA and Rac1 in 293 cells expressing either EphA1-WT or EphA1-P460L, stimulated with either pre-clustered ephrin-A1–Fc (1 μg/ml) or Fc (Control) for 15 minutes. (B,D) Densitometric quantification of (A) active Rac1 showing lower levels of active Rac1 in EphA1-P460L compared EphA1-WT-expressing 293 cells and (B) active RhoA showing higher levels of active RhoA in EphA1-P460L compared EphA1-WT-expressing 293 cells; data presented as the mean ± s.d. (N=4 independent experiments; two-way ANOVA and Bonferroni post-hoc analysis).

We also observed a concomitant increase in active RhoA-GTP levels in EphA1-P460L-expressing cells under basal (non-stimulated) conditions relative to EphA1-WT-expressing cells (Figure 6C, D). A similar increase in RhoA activity was seen in ligand-stimulated EphA1-WT-expressing cells again confirming that the kinase activity of EphA1 is required for regulating RhoA activity. Interestingly, unlike Rac1, we did observe a further increase in RhoA activity in EphA1-P460L-expressing cells, suggesting perhaps a secondary regulatory mechanism for RhoA activity by EphA1.

To confirm the involvement of RhoA and Rac1 in the spreading defect of EphA1-P460L-expressing cells, we set out to determine whether the phenotype could be rescued by restoring RhoA and Rac1 signaling to WT levels. To block RhoA activity in EphA1-P460L-expressing cells, we treated the cells with a specific inhibitor (Y-27632) against its downstream effector, Rho-associated kinase family proteins, ROCKI and ROCKII [39, 40] upon replating on fibronectin-coated plates. Cell were once again analyzed 30 minutes post-plating (Figure 7A-C). We found that blocking RhoA signaling in EphA1-P460L-expressing cells completely abrogated the spreading defect (Figure 7D, E). Next, we examined if restoring Rac1 activity could also rescue cell spreading phenotype. To accomplish this, we transiently co-transfected stably expressing EphA1-P460L cells with a constitutively active form of Rac1, Rac1-Q61L (Rac1-CA) [41] and an RFP-expressing construct for easy identification of transfected cells (Figure 8B, D). As performed above, cells were detached and replated on fibronectin-coated plates and analyzed after 30 minutes. We found that significantly more Rac1-CA expressing EphA1-P460L cells were able to spread normally, compared to non-transfected cells (Figure 8E, F). These data are consistent with our hypothesis that the P460L substitution on EphA1 alters receptor signaling and function, resulting in disrupted downstream RhoA and Rac1 signaling.

**Figure 7.**
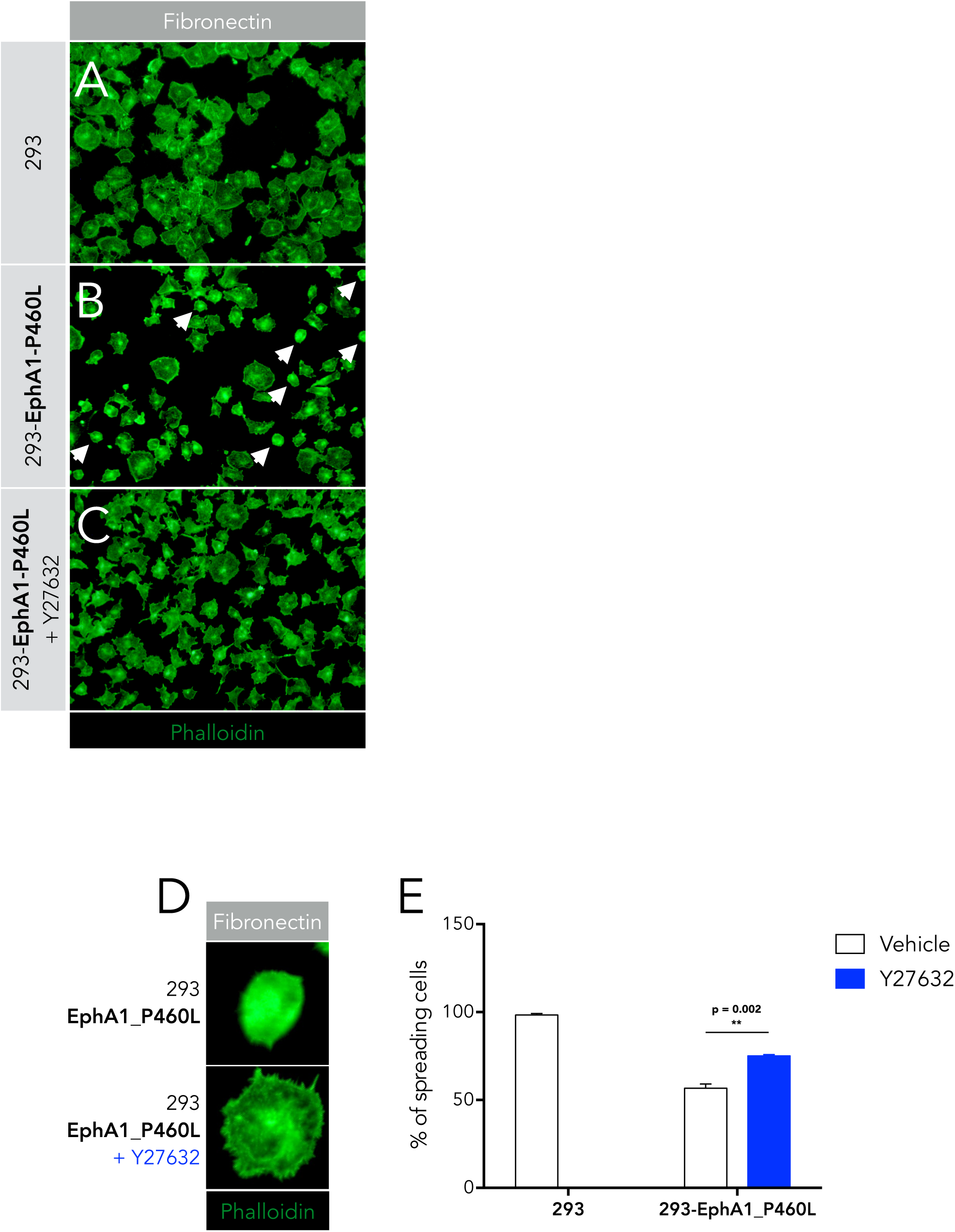
Delay in cell spreading by EphA1-P460L is dependent on RhoA activity. (A-D) Representative images of 293 cells stably expressing EphA1-P460L treated with either ROCK inhibitor Y-27632 (10 μM) or DMSO (control) and plated on FN-coated (1 μg/ml) coverslips for 30 minutes and stained with Alexa Fluor-488-conjugated Phalloidin (white arrows indicate round non-spreading cells). (E) Quantification of spreading cells expressed as the percentage of the total number of cells showing a higher percentage of spreading in EphA1-P460L expressing cells treated with Y-27632. Data presented as the mean ± s.d. (10 randomly selected high-power fields from N=3 independent experiments; two-tailed unpaired t-test).

**Figure 8.**
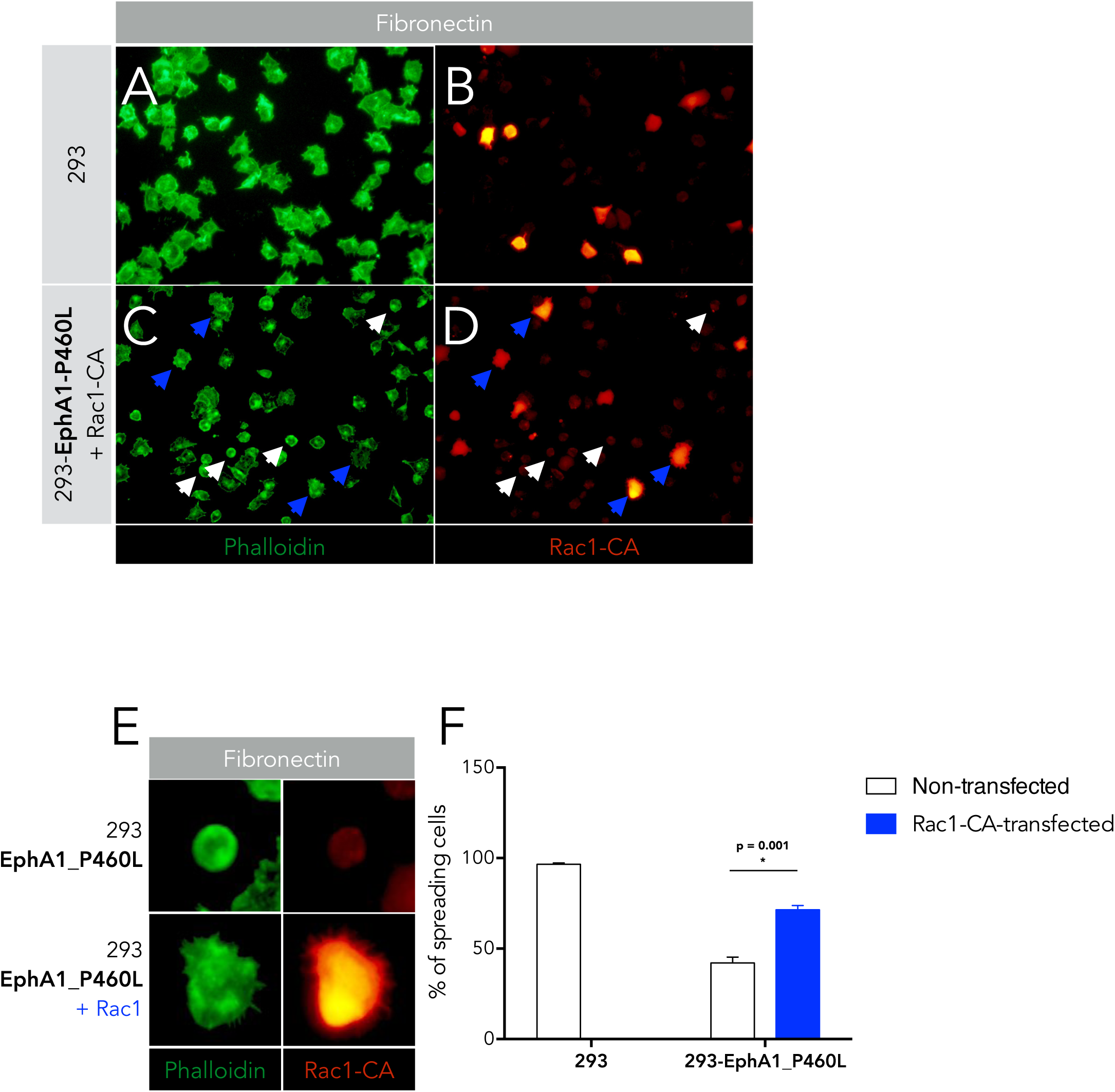
Delay in cell spreading by EphA1-P460L is rescued by restoring Rac1 activity. (A-E) Representative images of control 293 and cells stably expressing EphA1-P460L transiently transfected with constitutively active Rac1-Q61L and RFP and plated on FN-coated (1 μg/ml) coverslips for 30 minutes and stained with Alexa Fluor-488-conjugated Phalloidin (white arrows indicate round non-spreading and non-transfected cells; blue arrows indicate spreading and transfected cells). (E) Quantification of spreading cells expressed as the percentage of the total number of cells showing a higher percentage of spreading in EphA1-P460L expressing cells transfected with Rac-1-CA. Data presented as the mean ± s.d. (10 randomly selected high-power fields from N=3 independent experiments; two-tailed unpaired t-test).

## DISCUSSION

Characterization of AD-associated P460L variant of the Eph receptor EphA1 has never been reported. Here we have demonstrated the biological effects of the P460L amino acid substitution in EphA1 utilizing a biophysical, biochemical, and functional approach.

Our predicted structure modeling and analysis of EphA1-P460L suggests that this substitution potentially enhances anchoring of the EphA1 ectodomain deeper into the lipid bilayer through enhanced hydrophobic interactions with some of the components of lipid bilayers. This is supported by the fact that EphA1-P460L appears to interact more strongly with several components in the lipid bilayer. Lipid microdomains play an important role in finetuning the activity of many transmembrane receptors by positively or negatively affecting clustering and signal transduction [42]. The more hydrophobic amino acid, (leucine from proline) could cause the EphA1 variant to partially embed into the lipid bilayer and therefore restrict its diffusion. At low receptor concentrations, pre-clustered ligands are needed to promote initiate receptor clustering. However, above a certain threshold, free-moving EphR can cluster independently of ligand binding, through ectodomain interactions. Restricting movement could create “nucleation sites” which can recruit more receptors to the area and strengthen intercellular communication. This is further supported by the fact that our model shows the EphA1 P460L substitution to be located in the fibronectin type III repeats (FN2) proximal to the cell membrane, which has previously been reported to play an important role in helping ligand-independent [43].

Predictions based on our biophysical approach were confirmed biochemically. Immunoprecipitation analyses suggest that EphA1-P460L exists primarily as pre-formed clusters at the plasma membrane, with a lower concentration of monomers when compared to EphA1-WT. Consistent with this, immunoprecipitated EphA1-P460L were found to be significantly more tyrosine-phosphorylated than EphA1-WT, indicative of receptor auto-activation. In fact, the addition of pre-clustered ephrin-A1 ligand did not further increase the pool of tyrosine-phosphorylated EphA1-P460L, suggesting that the majority of EphA1-P460L may be phosphorylated. Activity of the receptor was confirmed by looking at several downstream targets of EphA1 signaling, including ERK1/2 and RhoA and Rac1. Together, these results suggest that EphA1-P460L may result in a constitutively active receptor.

Finally, functional assays revealed that EphA1-P460L significant delayed cell spreading through the aberrant Rho-GTPase signaling. This is supported by the observation that the spreading defect was rescued by correcting the RhoA/Rac1 dysregulation. Considering all our evidence, we propose a working model of the effect of the EphA1-P460 substitution on signaling and function, whereby constitutively EphA1-P460L leads to dysregulated RhoA/Rac1 signaling (Figure 9).

**Figure 9.**
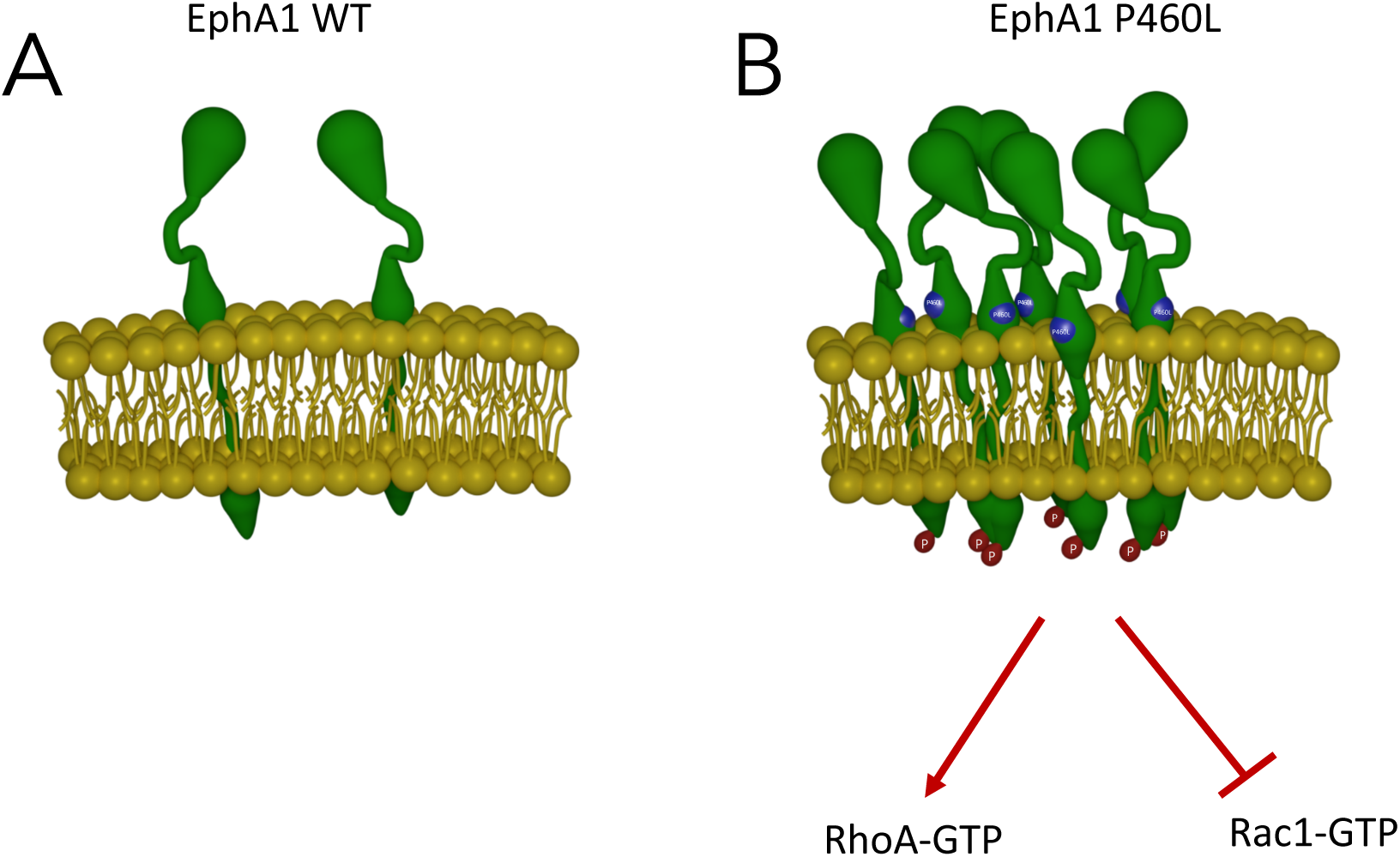
Working model of the effects of the P460L amino acid substitution on EphA1 function. Substitution of the more hydrophobic leucine residue enhances anchoring of the EphA1 ectodomain deeper into the lipid bilayer favoring the formation of receptor clusters and constitutive activation of EphA1 signaling independently of ligand binding. Constitutive activation of EphA1 signaling disrupts downstream targets RhoA and Rac1.

We propose that EphA1-P460L could affect susceptibility to LOAD through the dysregulation of these important pathways. Rho-GTPases are involved in a wide range of cellular processes such as mitosis, cell migration, adhesion and morphogenesis [44]. But it is their role in regulating dendritic spine morphology and synaptic function [45] and microglia function [46, 47] that makes these results particularly interesting. Several lines of evidence have shown that RhoA and Rac1 signaling is impaired in AD, potentially leading to synaptic dysfunction [28], tau hyperphosphorylation [48] or microglia dysfunction [49, 50]. As a result, Rho-GTPases have emerged as a potential important therapeutic targets [51].

Further investigation is needed to further elucidate not only how EphA1 variants influences AD pathogenesis but the role of EphA1 in neurodevelopment and neurodegeneration in general. This could provide further insight into how genetic risks factors act together to disrupt multiple signaling nodes, which may reveal novel targets.

## MATERIALS AND METHODS

### Materials

All chemicals used were of the highest grade available. Antibodies used in this study include the following: mouse anti-RhoA 67B9 (Cell Signaling, 1:2000), rabbit anti-Rac1 (cell signaling, 1:2000), mouse anti Phospho-Tyrosine P-Tyr-100 (Cell Signaling, 1:2000), rabbit anti Flag (Cell Signaling, 1:1000), mouse anti Myc-Tag 9B11 (Cell Signaling, 1:1000), rabbit anti-GAPDH (Cell Signaling, 1:2000). Reagents used in the study include: Glutamax (ThermoFisher Scientific), Fibronectin (FN) poweder (Sigma Aldrich), Dubelcco’s modified Eagle’s medium (DMEM) high glucose (ThermoFisher Scientific), fetal bovine serum (FBS, ThermoFisher Scientific), Complete Mini protease inhibitor cocktail (Roche Applied Science), Halt protease and phosphatase inhibitor (ThermoFisher Scientific), TRIzol reagent (Ambion), anhydrous dimethyl sulfoxide (DMSO, Sigma Aldrich), methylene chloride (Sigma Aldrich), phosphate-buffered saline pH 7.4 (PBS, ThermoFisher Scientific), tris-buffered saline pH 7.4 (TBS, ThermoFisher Scientific), Superblock (TBS) blocking buffer (ThermoFisher Scientific), paraformaldehyde 16% (PFA, ThermoFisher Scientific), Triton X-100 (ThermoFisher Scientific), radioimmunoprecipitation assay (RIPA) buffer (ThermoFisher Scientific), mammalian protein extraction reagent (M-PER, ThermoFisher Scientific), and DNase/RNase-free water (ThermoFisher Scientific). Extraction buffer (EB) recipe: 20 mM [(4-[2-hydroxyethyl]−1-piperazineethanesulfonic acid]) (HEPES)], pH 7.4, 100 mM NaCl, 20 mM NaF, 1 % Triton X-100, 1 mM sodium orthovanadate, 5 mM EDTA.

#### Sequence alignment

Sequence alignment of EphA1 and EphA2 receptors was carried out with T-coffee [52]. Domain boundaries were extracted from Uniprot [53].

#### Modeling of the wild-type and P460L Fibronectin domain FN2

Atomic modeling of the Fibronectin domain III-2 (residues 441-535) was carried out with I-TASSER [54]. We selected the atomic model with the highest C-value (0.12) and minimized it following a multi-step minimization protocol using the CHARMM forcefield and the conjugate gradient algorithm implemented in NAMD [55, 56]. First, hydrogen atoms were minimized for 3,000 steps, while heavy atoms were spatially constrained using a force constant of 10 Kcal/mol. Second, side-chain atoms were minimized for two rounds of 10,000 steps, while backbone atoms were spatially constrained using a force constant of 10 Kcal/mol and 2 Kcal/mol, respectively. Finally, all atoms were minimized for 200,000 steps without spatial constrains. P460L mutation was modelled on the minimized atomic model of the Fibronectin domain III-2 using Scap [57, 58]. Following this, the P460L atomic model was subjected to an additional round of minimization of the side-chains for 15,000 steps, while the backbone atoms were spatially constrained using a force constant of 10 Kcal/mol. The crystal structure of the EPHA2 FN2 was extracted from PDB 2×10 [34] and subjected to the same minimization protocol described above. Structural superposition of EPHA1 FN2 atomic model (wild-type and P460L mutant) and the EPHA2 FN2 anchored to the membrane [31] was performed with SKA [59].

### Electrostatic potential calculations

Electrostatic potentials were computed with the finite difference Poisson-Boltzmann (FDPB) method [60], implemented in Delphi [61]. Atomic charges and radii were extracted from the CHARMM forcefield [62]. The dielectric constant of the protein interior and the solvent were set to 2 and 80, respectively [62]. The ion exclusion parameter was set to 2 and the ionic strength to 145 mM. Electrostatic calculations were carried out using a lattice with 2.2 grids per Å and a series of focusing runs of increasing percentage fill (perfil) was performed from 20% to 90%. Calculations were iterated until they reached convergence, defined as the point at which the final maximum energy change is less than 10-4kTe-1. Visualization of electrostatic surfaces was carried out with Pymol.

### Cell culture and transfection

HEK 293 cells were maintained Dulbecco’s modified Eagle’s medium (DMEM) supplemented with 10% fetal bovine serum and 1% L-glutamine (check). The mammalian expression vectors for EphA1 WT and EphA1 P460L were transfected into HEK cells using Fugene6 transfection kit (Promega). Expression of EphA1 WT or EphA1 P460L was confirmed by immunoblotting with anti-Flag or anti-Myc antibody.

### qPCR

Cells were lysed and total RNA was extracted using TRIzol reagent (Ambion) following the manufacturer’s protocol. RNA concentration and purity were assessed by measuring the optical density at 260 and 280 nm with a NanoDrop (Thermo Scientific). cDNA was synthesized using the first-strand cDNA synthesis kit (Origene) with 1 μg of total RNA, following the manufacturer’s instructions. Quantitative real-time PCR was performed using FastStart SYBR Green Master Mix (Roche) and an Eppendorf Realplex Mastercyler with the following cycle settings: 1 cycle at 95°C for 10 min and 40 cycles of amplification, 95°C for 15 s, 58–60°C for 30–60 s, 72°C for 30–60 s. The amounts of EphA1 mRNA were quantified and normalized to GAPDH mRNA using the following primer pairs: EphA1 forward 5′-CATGGGGCATGCAGAGTCA-3′ and reverse 5′-CTGGCCCATGATAGTTGCCT-3′; The threshold cycles were determined and normalized to the threshold cycles of GAPDH.

### Immunoprecipitation and immunoblotting

Cells were lysed in RIPA buffer (Pierce) supplemented with phosphatase inhibitor and protease inhibitor for 20 minutes at 4°C. Lysates were spun down at 15,000g for 15 minutes at 4°C. Immunoprecipitation was carried out using antibodies at 1ug/mg of total protein at 4°C for at least for 1 hour. Immune complexes were collected using Dynabeads M-280 Streptavidin (ThermoFisher) using the manufacturer’s protocol. Briefly, collected beads were washed with immunoprecipitation washing buffer and proteins eluted in provided buffer. The samples were boiled in reducing SDS sample buffer. Proteins were transferred to nitrocellulose membrane (Invitrogen) and probed with the indicated antibodies.

### Cell spreading assays

HEK293 cells were plated on fibronectin-coated (1 μg/ml) coverslips with or without immobilized ephrin-A1–Fc (1 μg/ml) in adhesion medium (serum-free DMEM containing 0.1% BSA) and allowed to spread for 30 minutes at 37°C. Cells were observed under an Olympus IX70 fluorescent microscope equipped with digital camera and recorded. Images were analyzed by Image J to determine circularity. Non-spreading cells were defined as round cells (circularity=0.8-1), whereas spreading cells were defined as those with extended processes as described (Fukuda et al., 2003; Zhang et al., 2002). The percentage of cells with spreading morphology was quantified by analyzing at least 300 cells from ten randomly selected fields. For experiments with Rac1-Q61L, cells were transiently transfected with pRK5-myc-Rac1-Q61L (Addgene plasmid # 12983; http://n2t.net/addgene:12983; RRID:Addgene_12983) with Lipofectamine (ThermoFisher) following manufacturer’s protocol for 24 hours prior to experiments.

### Rho-GTPase activity assay

Rho-GTPase pull-down assays were performed following the manufacturer’s protocol (Cytoskeleton, Inc.). Briefly, cells were washed once with ice-cold PBS and lysed with RIPA/M-PER buffer (1:1; Thermo Fisher). Lysates were incubated on ice for 5 minutes, and centrifuged (12,000 g) for 5 minutes. 30 μl was used for input and the remaining lysates were incubated with Rhotekin-RBD (RhoA) or PAK-RBD (Rac1) protein beads (25μl) by rotating for 1 hour at 4°C. Beads were washed with ice-cold PBS (500 μl, 3x) and collected, and bound proteins were eluted with 1x sample buffer (30 μl). Input and eluted samples were then analyzed by gel electrophoresis and western blotting using either a RhoA or Rac1-specific antibody.

### Statistics

All statistical analyses were performed using GraphPad Prism software. Statistical significance was determined by Student’s t-test or one- or two-way analysis of variance (ANOVA), where appropriate, with a significance threshold of p < 0.05.

## Acknowledgements

This work was supported in part by grants from the Columbia University Alzheimer’s Disease Research Center (ADRC), from the Taub Institute for Research on Alzheimer’s disease and the\ Aging Brain, and from the Alzheimer Association.

## Contributions

R.L., G.L. and Y.K. conceived and designed the study. R.L., Y.K., G.L. and H.P. performed the experiments and analyzed data. R.L. and Y.K. wrote the manuscript. All authors discussed the results and contributed to the manuscript.

## Corresponding author

Correspondence to Roger Lefort.

## Ethics declarations

Competing interests

The authors declare no competing interests.

