## Supplementary Figures for "Alzheimer’s disease-associated P460L mutation in ephrin receptor type A1 (EphA1) leads to dysregulated Rho-GTPase signaling"

A

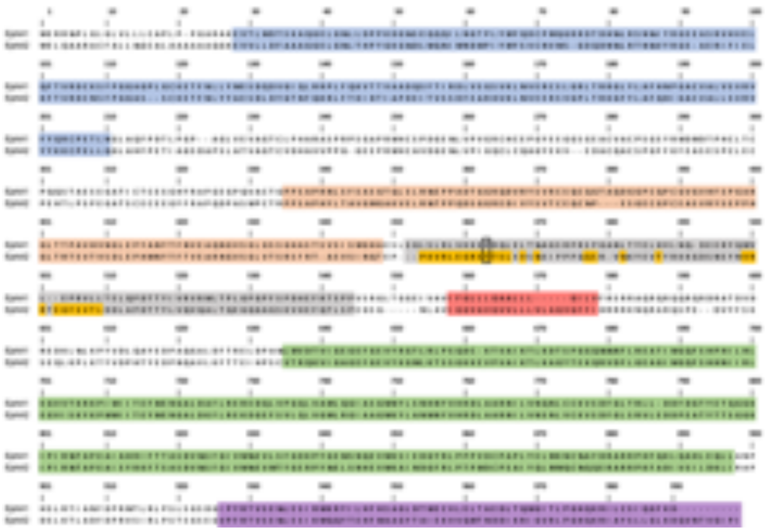

B

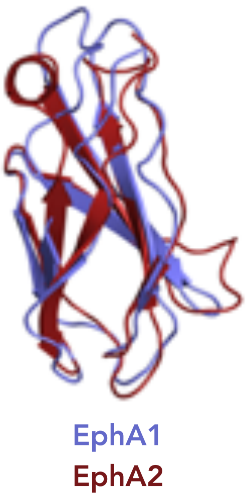


**Figure S1. Sequence alignment and domain organization of EphA1 and EphA2 receptors.** (A) Ligand binding domain (blue), fibronectin domain III 1 (salmon), fibronectin domain III 2 (grey), transmembrane domain (red), protein kinase domain (green), SAM domain (purple). Domain boundaries were retrieved from the Uniprot database (REF). EphA2 FN2 residues located within 4 Å of the lipid bilayer, as modelled by molecular dynamics (Chavent M., et. Al, Structure 2016), are shown in yellow. Alzheimer-associated EphA1 P460L mutation and the corresponding EphA2 aligned residue are shown in a box.

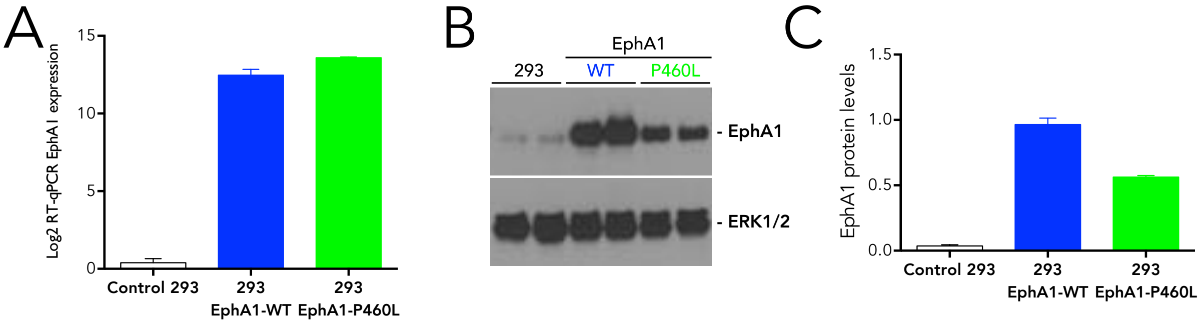


**Figure S2. Characterization of stable HEK293 cells lines overexpression EphA1-WT and EphA1-P460L |** (A) Quantitative-PCR analysis of EphA1 mRNA of stable 293 cells. (B,C) Representative Western blot and quantification showing protein expression of EphA1-WT and EphA1-P460L in stable 293 cells. Data presented as mean +/- s.d. values of N=3 independent cultures.

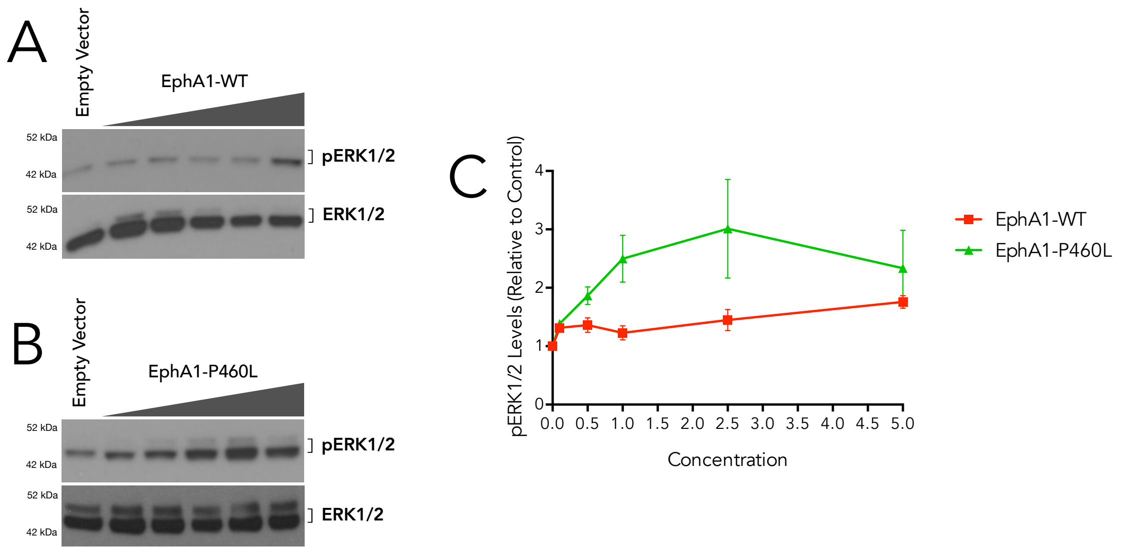


**Figure S3: Enhanced activation of downstream pathways of EphA1-P460L |** Amino acid substitution P460L in EphA1 results in enhanced phosphorylation of one of its downstream targets ERK1/2: EphA1 (WT and P460L) was overexpressed in 293 cells at increasing concentrations. (A-B) Representative immunoblot (phospho-ERK1/2) showing higher levels of phosphorylated ERK in EphA1-P460 at much lower doses then EphA1-WT consistent with a gain-of-function mutation. (C) Densitometric quantification of (A,B) ERK1/2 showing higher levels of ERK1/2 in cells expression EphA1-P460L compared to EphA1-WT. Data presented as the mean ± s.d. (N=4 independent experiments).

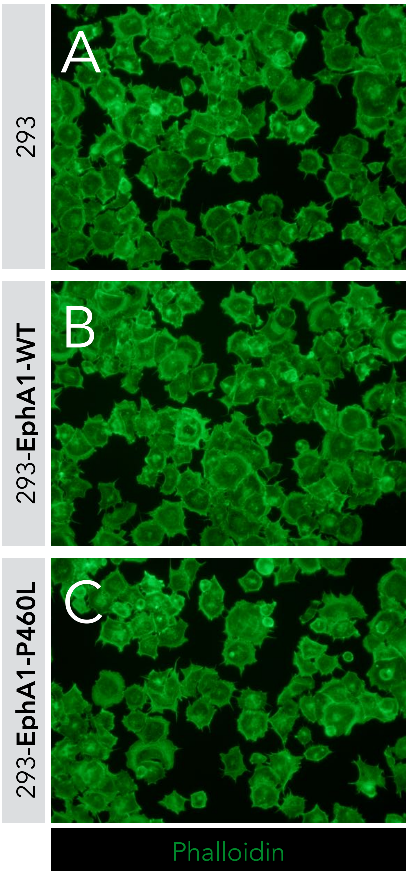


**Figure S4: Cell-spreading is delayed in EphA1-P460L cells |** Representative images of wild-type 293 (A) and stably expressing EphA1-WT (B) or EphA1-P460L (C) plated on fibronectin (FN)-coated (1 μg/ml) coverslips for 30 minutes and stained with Alexa Fluor-488-conjugated Phalloidin.
